# Functional analysis of deoxyhexose sugar utilization in *Escherichia coli* reveals fermentative metabolism under aerobic conditions

**DOI:** 10.1101/2021.04.12.439581

**Authors:** Pierre Millard, Julien Pérochon, Fabien Letisse

## Abstract

L-rhamnose and L-fucose are the two main 6-deoxyhexoses *Escherichia coli* can use as carbon and energy sources. Deoxyhexose metabolism leads to the formation of lactaldehyde whose fate depends on oxygen availability. Under anaerobic conditions, lactaldehyde is reduced to 1,2-propanediol whereas under aerobic condition, it should be oxidised into lactate and then channelled into the central metabolism. However, although this all-or-nothing view is accepted in the literature, it seems overly simplistic since propanediol is also reported to be present in the culture medium during aerobic growth on L-fucose. To clarify the functioning of 6-deoxyhexose sugar metabolism, a quantitative metabolic analysis was performed to determine extra- and intracellular fluxes in *E. coli* K-12 MG1655 (a laboratory strain) and in *E. coli* Nissle 1917 (a human commensal strain) during anaerobic and aerobic growth on L-rhamnose and L-fucose. As expected, lactaldehyde is fully reduced to 1,2-propanediol in anoxic conditions allowing complete reoxidation of the NADH produced by glyceraldehyde-3-phosphate-dehydrogenase. We also found that net ATP synthesis is ensured by acetate production. More surprisingly, lactaldehyde is also primarily reduced into 1,2-propanediol under aerobic conditions. For growth on L-fucose, ^13^C-metabolic flux analysis revealed a large excess of available energy, highlighting the need to better characterize ATP utilization processes. The probiotic *E. coli* Nissle 1917 strain exhibits similar metabolic traits, indicating that they are not the result of the K-12 strain’s prolonged laboratory use.

**IMPORTANCE:** *E. coli*’s ability to survive, grow and colonize the gastrointestinal tract stems from its use of partially digested food and hydrolysed glycosylated proteins (mucins) from the intestinal mucus layer as substrates. These include L-fucose and L-rhamnose, two 6-deoxyhexose sugars, whose catabolic pathways have been established by genetic and biochemical studies. However, the functioning of these pathways has only partially been elucidated. Our quantitative metabolic analysis provides a comprehensive picture of 6-deoxyhexose sugar metabolism in *E. coli* under anaerobic and aerobic conditions. We found that 1,2-propanediol is a major by-product under both conditions, revealing the key role of fermentative pathways in 6-deoxyhexose sugar metabolism. This metabolic trait is shared by both *E. coli* strains studied here, a laboratory strain and a probiotic strain. Our findings add to our understanding of *E. coli*’s metabolism and of its functioning in the bacterium’s natural environment.

## INTRODUCTION

Rhamnose and fucose are the two main naturally occurring 6-deoxyhexose sugars, found mainly in their L enantiomeric forms. L-Rhamnose, formed in bacteria and plants but not in humans, is mainly found in heteropolymers such as hemicellulose, the second largest component of lignocellulose (1). In contrast, L-fucose occurs mostly in prokaryotes and eukaryotes in glycoconjugates, such as mucin in intestinal mucus (2). L-fucose frequently occupies the terminal position in common mucin glycoproteins, lining the epithelium with L-fucose moieties. L-rhamnose and L-fucose are released by chemical or enzymatic hydrolysis from dietary fibres or fucosylated oligosaccharides in the gastrointestinal tract, where they can then be used by many bacteria to support growth and intestinal colonization (3, 4). *Escherichia coli* is the predominant facultative anaerobe in human gastrointestinal tract ecosystems and is commensal in the gut, colonizing it lifelong from an early age (5). *E. coli’s* ability to colonize the intestine results mainly from its capacity to draw nutrients from mucus, notably L-fucose, which appears to be involved in its maintenance (6) as commensal *E. coli* strains that lack the ability to metabolize L-fucose show defective colonization (7).

The earliest studies of *E. coli* metabolism of L-fucose and L-rhamnose date back to the 1970s (8). The first steps of L-fucose and L-rhamnose catabolism lead to the formation of dihydroxyacetone-phosphate (DHAP, C1 to C3 fragment) and S-lactaldehyde (C4 to C6 fragment) (Figure 1). DHAP is an intermediate of the central metabolism that can participate in both gluconeogenic and glycolytic processes. In contrast, S-lactaldehyde’s fate is less variegated and depends on oxygen availability. Aerobically, S-lactaldehyde is converted through two successive steps of oxidation into pyruvate, the first, catalysed by a (NADH)-linked aldehyde dehydrogenase (AldA) leading to the formation of S-lactate, and the second, catalysed by a FMN-dependent membrane-associated S-lactate dehydrogenase (LldD). Anaerobically, S-lactaldehyde is reduced by S-1,2-propanediol oxydoreductase (FucO) into S-1,2-propanediol as a terminal electron acceptor, which is excreted into the environment. Theses enzymes are encoded by genes organized in operons, the *fucPIKUR* and *fucAO* operons for the fucose regulon (9) and the *rhaBAD* and *RhaT* operons for the rhamnose regulon (10), which lacks its own propanediol oxidoreductase. The expression of *fucO* on rhamnose must therefore result from cross induction of the fucose enzymes through an incompletely understood mechanism (9).

**Figure 1.**
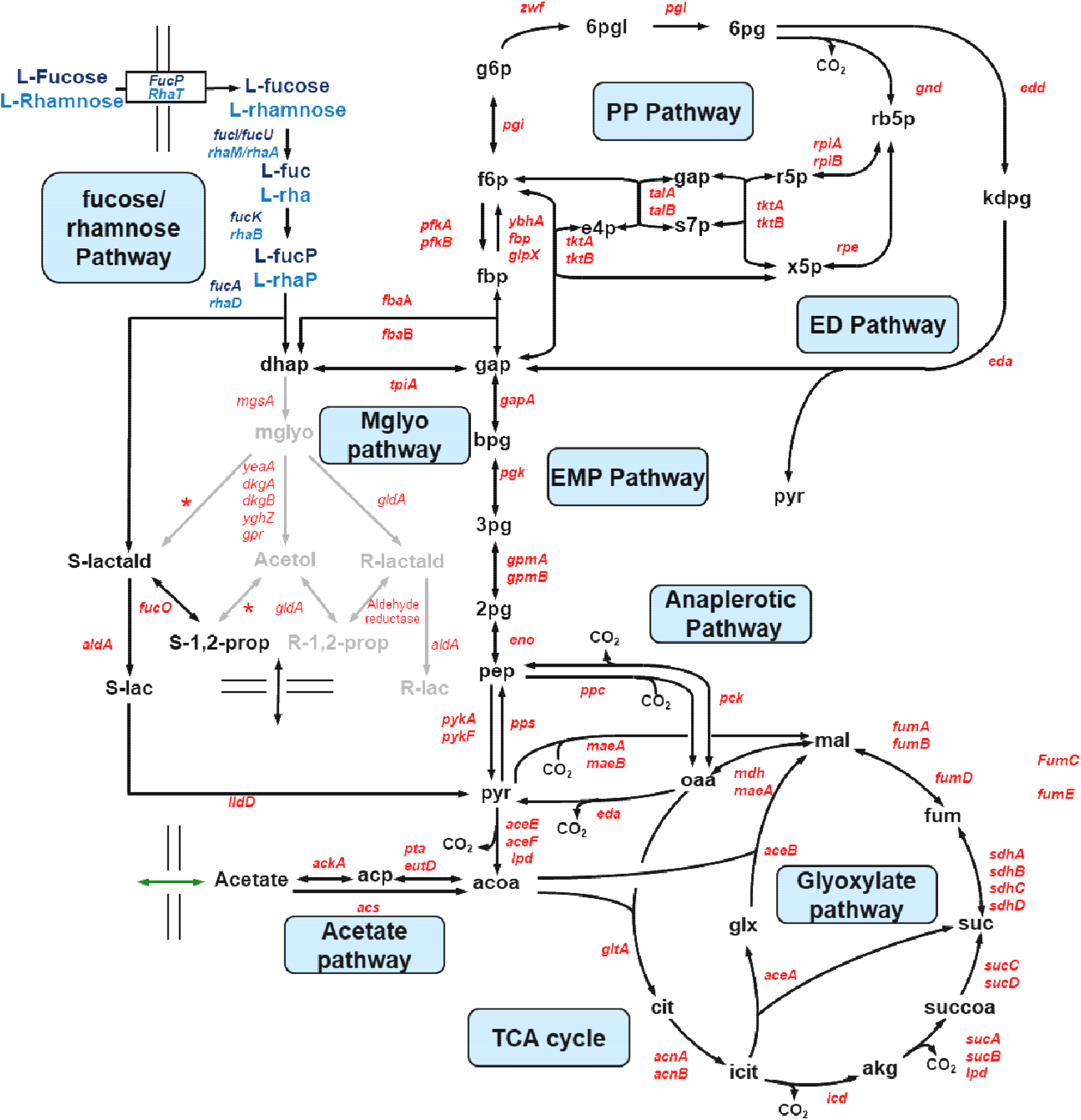
Metabolic network of 6-deoxyhexose catabolism in *Escherichia coli*. The genes encoding the various enzymatic steps are shown in italics. The genes and metabolites specific to fucose and rhamnose metabolism are shown in dark blue and medium blue respectively. *, suspected reactions with no identified gene(s) (21). Methylgloxal metabolism is shown in grey. Pathway abbreviation: PP pathway, pentose-phosphate pathway; ED pathway, Entner-Doudoroff pathway; EMP pathway, Embden-Meyerhof-Parnas pathway (glycolysis); Mglyo pathway, methylglyoxal pathway; TCA cycle, tricarboxylic acid cycle. Metabolite abbreviations: L-fuc, L-fuculose; L-rha, L-rhamnulose; L-fucP, L-fuculose-1-phosphate; L-rhaP, L-rhamnulose-1-phosphate; S(R)-lactald, S(R)-lactaledyde; S(R)-lact, S(R)-lactate; S(R)-1,2-prop, S(R)-1,2-propanediol; Mglyo, methylglyoxal; g6p, glucose-6-phosphate; f6p, fructose-6-phosphate; fbp, fructose-1,6-biphosphate; dhap, dihydroxyacetone phosphate; gap, glyceraldehyde 3-phosphate; bpg, 1,3-biphosphoglycerate; 3pg, 3-phosphoglycerate; 2pg, 2-phophoglycerate; pep, phosphoenolpyruvate; pyr, pyruvate; 6pgl, 6-phosphoglucono-δ-lactone; 6pg, 6-phophogluconate; Rb5P, ribulose-5-phosphate; r5p, ribose-5-phosphate; x5p, xylulose-5-phosphate; s7p, sedoheptulose-7-phosphate; e4p, erythrose-4-phosphate; acoa, acetyl coenzyme A; acp, acetyl phosphate; cit, citrate; icit, isocitrate; glx, glyoxylate; akg, α-ketoglutarate; succoa, succinyl-coenzyme A; suc, succinate; fum, fumarate; mal, malate; oaa, oxaloacetate.

While the distinct anaerobic/aerobic fates of S-lactaldehyde are well described in the literature, albeit mainly through genetic and biochemical studies (11–13), little is known about the actual distribution of S-lactaldehyde - and to a larger extent, DHAP - within central metabolic pathways. Zhu and Lin (13) found for example that S-lactaldehyde is partially reduced into 1,2-propanediol despite the presence of oxygen, meaning that the oxidation and reduction of S-lactaldehyde are not all-or-nothing processes exclusively determined by the presence or absence of oxygen. To date, studies of L-fucose and L-rhamnose metabolism in *E. coli* have provided only an extremely partial view of the operation of the central metabolism and no insight into the cellular redox and energy metabolisms when these compounds are the sole carbon and energy sources.

Therefore, with the aim of clarifying 6-deoxyhexose sugar metabolism in *E. coli*, we carried out a quantitative metabolic analysis involving exometabolome and metabolic flux analyses across the central metabolism, of *E. coli* strains (the model K-12 MG1665 strain and *E. coli* Nissle 1917) grown under controlled anaerobic and aerobic conditions on L-fucose and L-rhamnose. Nissle 1917 is a probiotic strain, classified in the B2 phylogenetic group (14), which is over-represented in the human microbiota in developed countries (15) and contains most pathogenic *E. coli* strains. These strains are efficient colonizers of the gut (14) and *E. coli* Nissle 1917 is therefore a representative strain whose metabolic traits, unlike those of K-12-like strains, have not been affected by prolonged laboratory use (16). Our results notably reveal an unexpected fermentative metabolism of L-fucose and L-rhamnose, characterized by the excretion of S-1,2-propanediol even in the presence of oxygen, and updates our current knowledge of 6-deoxyhexose sugar metabolism in *E. coli*.

## RESULTS SECTION

### Anaerobic growth of *E. coli* K-12 on 6-deoxysugars

We first sought to establish the detailed macrokinetics of *Escherichia coli* growing anaerobically in minimal medium containing 10 g.L^-1^ L-fucose or L-rhamnose as sole carbon source. The growth experiments were performed in bioreactors at 37°C, pH 7, under nitrogen atmosphere.

Growth of *E. coli* on fucose is accompanied by 1,2-propanediol (1,2-PDO), formate and acetate production (Figure 2A). These end-products represent nearly 88% (Cmol/Cmol) of the carbon balance, the rest being mainly biomass (∼5% Cmol/Cmol) and CO_2_ (not measured). The biomass yield is somewhat lower than expected under anaerobic conditions (Table 1). The growth rate (0.20 ± 0.01 h^-1^) is similar to the one reported by Boronat and Aguilar (17). Interestingly, only traces of lactate and ethanol are detected even though the latter is the main redox balancing end-product, when *E. coli* is grown anaerobically on glucose for example (18, 19). The 1,2-PDO yield is approximatively 1 mol/mol of fucose consumed. This means that lactaldehyde is fully converted into 1,2-PDO since the breakdown of 1 mole of fuculose-1-phosphate yields 1 mole of DHAP and 1 mole of lactaldehyde. The oxidation of NADH in the endergonic part of glycolysis from DHAP is thus fully effected by the reduction of lactaldehyde into 1,2-PDO (Figure 3), balancing the NADH fluxes. This explains why no ethanol is detected, but significant amounts of acetate are (0.76 ± 0.05 mol/mol of fucose consumed), with net synthesis of ATP, which in the absence of oxygen or other external electron acceptors can only be synthetized by substrate-level phosphorylation. The net synthesis of ATP was estimated to be 43.8 ± 2.0 mmol.(g_CDW_.h)^-1^, which is apparently sufficient to cover the demands for ATP in the biosynthesis of anabolic precursors, in growth- and non-growth-associated maintenance, and in fucose uptake (see below).

**Table 1:** Growth parameters of *E. coli* K-12 MG1655 cultivated aerobically and anaerobically on L-fucose or on L-rhamnose as sole carbon sources. The data shown are the specific rates for growth (μ_max_), substrate uptake (q_S_), 1,2-propanediol formation (q_PDO_), acetate formation (q_Ace_), and formate formation (q_For_), calculated from the data in Figure 2, and the yields of biomass (Y_X/S_), 1,2-propanediol (Y_PDO/S_), acetate (Y_Ace/S_) and formate (Y_For/S_).

| Parameters | L-fucose |  | L-rhamnose |  |
| --- | --- | --- | --- | --- |
|  | -O<sub>2</sub> | +O<sub>2</sub> | -O<sub>2</sub> | +O<sub>2</sub> |
| $\mu$ (h<sup>-1</sup>) | 0.20 ± 0.01 | 0.49 ± 0.01 | 0.27 ± 0.01 | 0.36 ± 0.01 |
| $q_s$ (mmol.(g<sub>CDW</sub>.h)<sup>-1</sup>) | 27.5 ± 0.7 | 15.3 ± 0.3 | 19.1 ± 0.1 | 10.0 ± 0.2 |
| $q_{\text{PDO}}$ (mmol.(g<sub>CDW</sub>.h)<sup>-1</sup>) | 29.4 ± 1.1 | 11.6 ± 0.1 | 16.7 ± 0.1 | 7.1 ± 0.1 |
| $q_{\text{Ace}}$ (mmol. (g<sub>CDW</sub>.h)<sup>-1</sup>) | 21.0 ± 1.3 | 4.5 ± 0.1 | 13.1 ± 0.1 | 0.7 ± 0.1 |
| $q_{\text{For}}$ (mmol. (g<sub>CDW</sub>.h)<sup>-1</sup>) | 18.3 ± 0.1 | ND* | 10.4 ± 0.1 | ND* |
| $q_{\text{CO}_2}$ (mmol. (g<sub>CDW</sub>.h)<sup>-1</sup>) | NM** | 17.8 ± 0.9 | NM** | 17.6 ± 0.4 |
| $Y_{X/S}$ (g<sub>X</sub>.mol<sup>-1</sup>) | 7.2 ± 0.2 | 31.7 ± 0.6 | 14.3 ± 0.1 | 35.8 ± 0.9 |
| $Y_{\text{PDO}}$ (mol.mol<sup>-1</sup>) | 1.07 ± 0.05 | 0.76 ± 0.02 | 0.87 ± 0.01 | 0.71 ± 0.02 |
| $Y_{\text{Ace}}$ (mol.mol<sup>-1</sup>) | 0.76 ± 0.05 | 0.30 ± 0.01 | 0.69 ± 0.01 | 0.07 ± 0.01 |
| $Y_{\text{For}}$ (mol.mol<sup>-1</sup>) | 0.68 ± 0.02 | ND* | 0.54 ± 0.01 | ND* |
| $Y_{\text{CO}_2}$ (mol.mol<sup>-1</sup>) | NM* | 1.17 ± 0.06 | NM* | 1.75 ± 0.05 |

\*: Not determined
\*\* : Not measured

**Figure 2.**
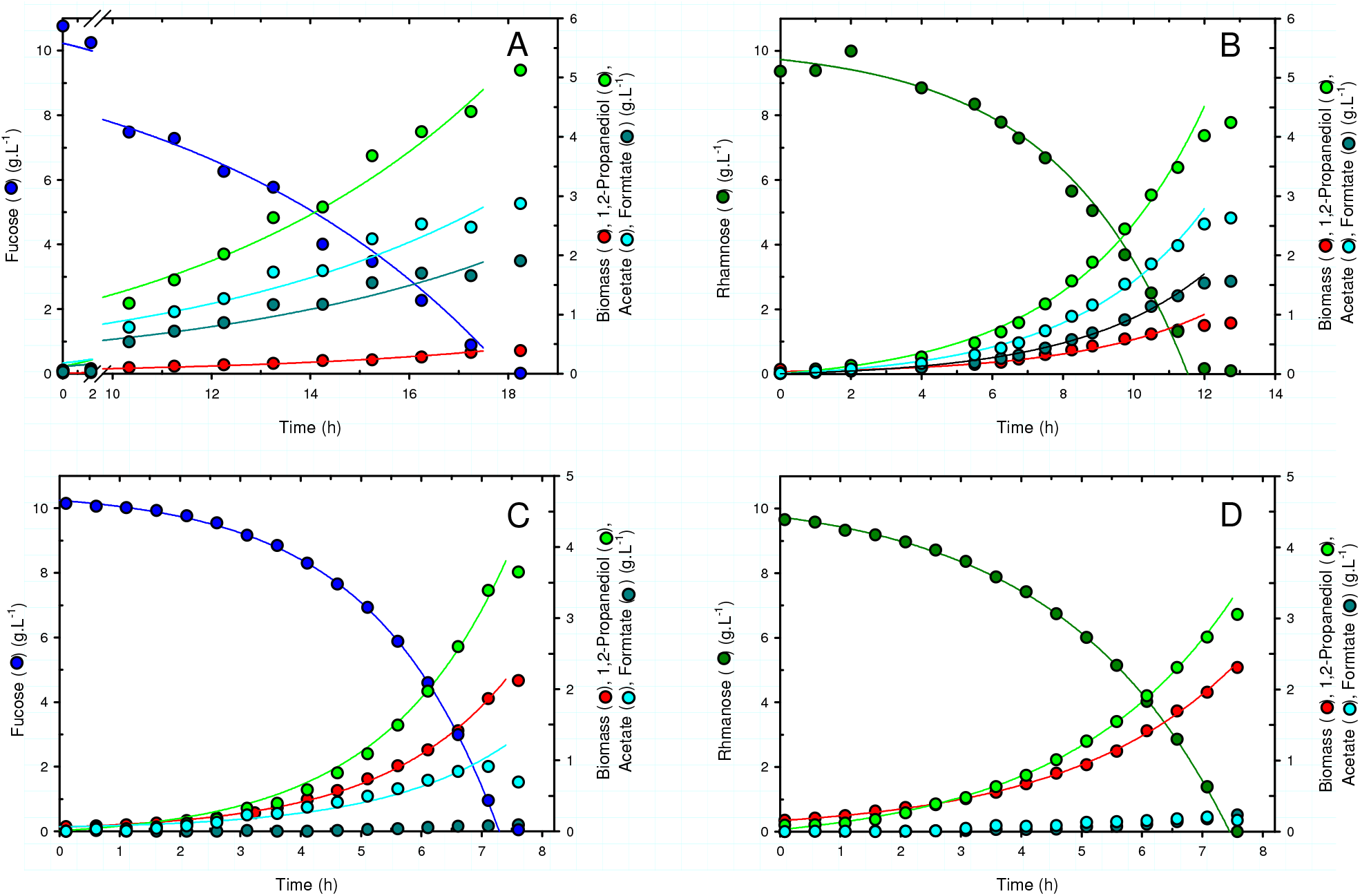
Growth profiles of *E. coli* K-12 MG1655 cultivated anaerobically on fucose (A) or rhamnose (B), and aerobically on fucose (C) or rhamnose (D). The cultures were grown in bioreactors on fucose or rhamnose as sole carbon and energy source, under nitrogen atmosphere for anaerobic cultures and with a dissolved oxygen tension above 30% for aerobic cultures. The solid lines are the best fits obtained with physiofit (43).

**Figure 3.**
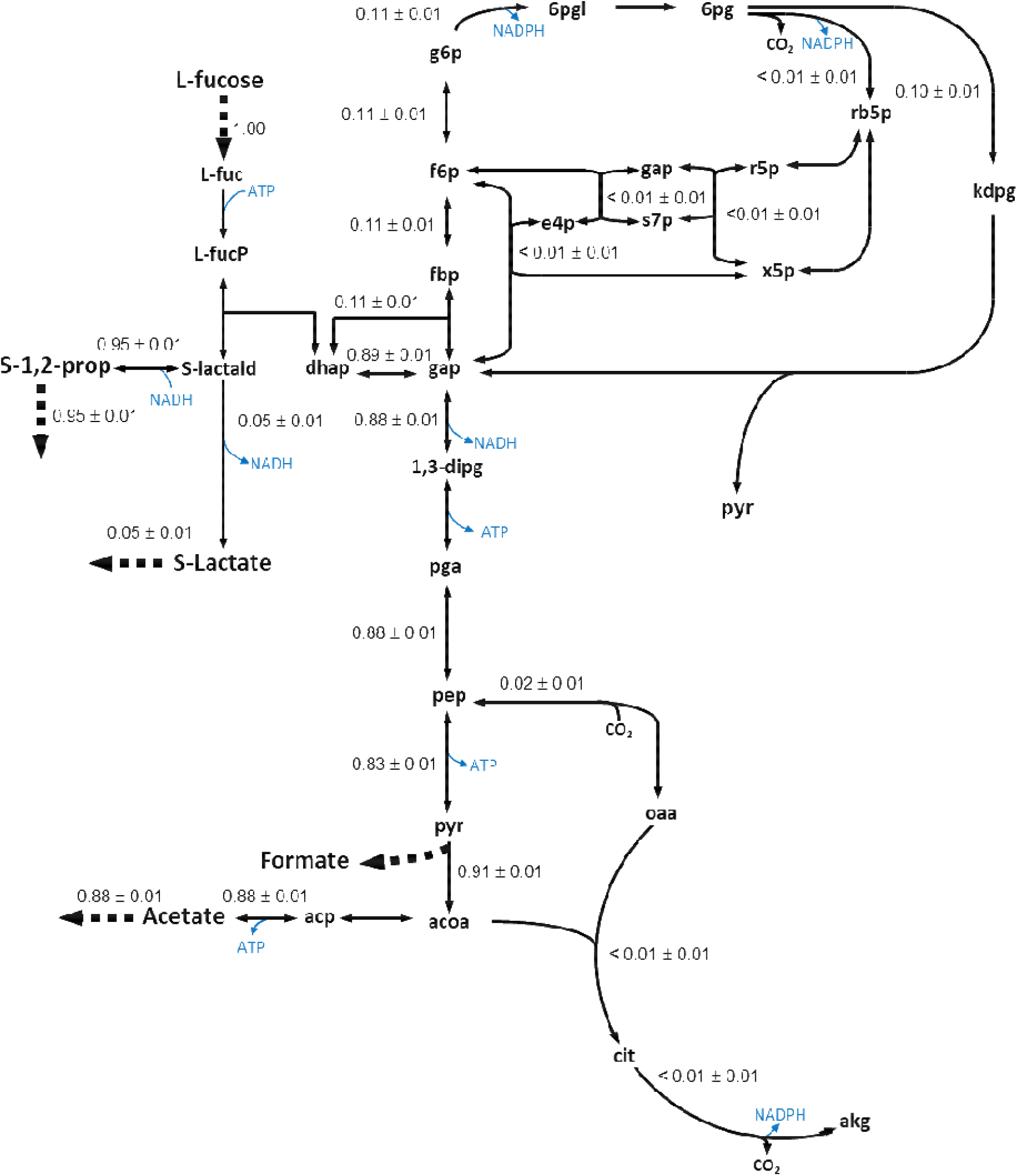
Flux distribution in the central metabolism of *Escherichia coli* K-12 MG1655 growing anaerobically on fucose. Fluxes are given as a molar percentage of the specific fucose uptake rate which was set to 1.

Although anaerobic growth on rhamnose appears to be significantly faster, the pattern of end-products excreted in culture broths of *E. coli* K-12 and the respective yields are similar to those determined for growth on fucose (Figure 2B - Table 1), indicating that the anaerobic metabolism of rhamnose is analogous to the one described for fucose.

### Aerobic growth of *E. coli* K-12 on 6-deoxyhexose sugars

In the same culture system under aerobic conditions (DOT>30%), *E. coli* grows exponentially on L-fucose at 0.49 h^-1^, 2.5-fold faster than under anaerobic conditions (Table 1). In keeping with what it is generally observed with other carbon sources, the specific fucose consumption rate (q_fuc_) under aerobic conditions is lower than under anaerobic conditions (15.3 ± 0.3 mmol.(g_CDW_.h)^-1^ *versus* 27.5 ± 0.7 mmol.(g_CDW_.h)^-1^). More importantly, we observed that *E. coli* grown aerobically on fucose produces large amounts of 1,2-PDO (Figure 2C), with a yield of approximately 0.76 mol 1,2-PDO/mol of fucose consumed. This was not expected since under these conditions, S-lactaldehyde is supposed to be converted into pyruvate *via* two successive oxidations catalyzed respectively by AldA and LldD (8, 9, 13) (Figure 1), such that the metabolic flux should be fully channeled towards the tricarboxylic acid cycle. Note however that the pioneering studies that established this all-or-nothing view of 6-deoxyhexose sugar metabolism also found significant production of 1,2-PDO in the presence of oxygen (8, 13).

Also produced in significant amounts alongside 1,2-PDO is acetate (about 10% Cmol/Cmol of fucose consumed), as observed in fast growing *E. coli* cells on glucose (20). However, the acetate production rate and yield are much lower than under anaerobic conditions. Apart from 1,2 PDO and acetate, no other by-products were detected in significant amounts.

*E. coli* K-12 grows similarly on L-rhamnose as on L-fucose (Figure 2D), but except for formate production, with substantially lower growth parameters (Table 1). Growth and rhamnose consumption rates (q_rha_) are diminished by approximatively 30% (the biomass yield is indeed similar on both substrates) without any significant effect on the production of 1,2-PDO, which accumulates in the culture medium with a similar molar yield as estimated on fucose. In contrast, acetate overflow is reduced and CO_2_ production increased.

The key results here are that *E. coli* K-12 produces 1,2-propanediol at high yield when grown on L-fucose and L-rhamnose under aerobic conditions. This represents a serious waste of carbon as up to 38% of the 6-deoxyhexose is converted into 1,2-PDO. 1,2-PDO could be formed directly from S-lactaldehyde *via* propanediol oxidoreductase (POR) encoded by the gene *fucO* in the fucose regulon, in a similar manner as under anaerobic conditions. Alternatively, 1,2-PDO could also be formed from DHAP *via* methylglyoxal synthase (MGS) and the enzymes involved in the detoxification of the resulting methylglyoxal, such as glycerol dehydrogenase (GldA) (Figure 1).

### 1,2-PDO is formed by propanediol oxidoreductase

One way to determine the extent to which each of these routes contributes to 1,2-PDO formation is to quantify the relative amounts of the two enantiomeric forms produced. Indeed, POR’s conversion of S-lactaldehyde into S-1,2-PDO is enantiomerically pure since it does not reduce R-lactaldehyde (17). In contrast, the 1,2-PDO synthesized from methylglyoxal, either *via* acetol or R-lactaldehyde, is mainly the R enantiomer (21). The ratio of 1,2-PDO enantiomers produced is therefore indicative of the relative contributions of the two metabolic routes involved in its synthesis.

The enantiomeric purity of the 1,2-PDO produced by *E. coli* K-12 grown on L-fucose and L-rhamnose was determined by liquid chromatography with a chiral column. Culture supernatants collected from the aerobic cultures gave a single peak at the retention time of S-1,2-PDO (Figure S1). Since no R-1,2-PDO was detected, this indicates that during aerobic growth on 6-deoxyhexose sugars the 1,2-PDO produced by *E. coli* K-12 is formed by POR from S-lactaldehyde.

### Intracellular carbon and energy fluxes on fucose

To obtain a quantitative understanding of the functioning of the metabolic network, we determined the intracellular flux distribution in *E. coli* during aerobic growth on L-fucose. We focused on L-fucose because the ^13^C-labeled forms required for this experiments are commercially available and relatively inexpensive, which is not the case for L-rhamnose. *E. coli* K-12 was therefore grown on a mixture of 80% 1-^13^C-L-fucose and of 20% U-^13^C-L-fucose as sole carbon source in a 1 L baffled shake-flask containing 80 mL of medium to reduce the amount of labelled fucose used.

The growth kinetics were similar to those observed in bioreactors, though the 1,2-PDO yield was slightly lower (Figure 4). The metabolic flux map was established from the growth rate, extracellular fluxes, and ^13^C-incorporation quantified in central metabolites and proteinogenic amino acids (22), using a detailed isotopic model of *E. coli*’s central carbon metabolism (23). A chi-square statistical test confirmed that the isotopic model fitted the data satisfactorily (p-value < 0.05), and this was also supported by the good correlation between measured and simulated data (R^2^ > 0.99, Figure S2).

**Figure 4.**
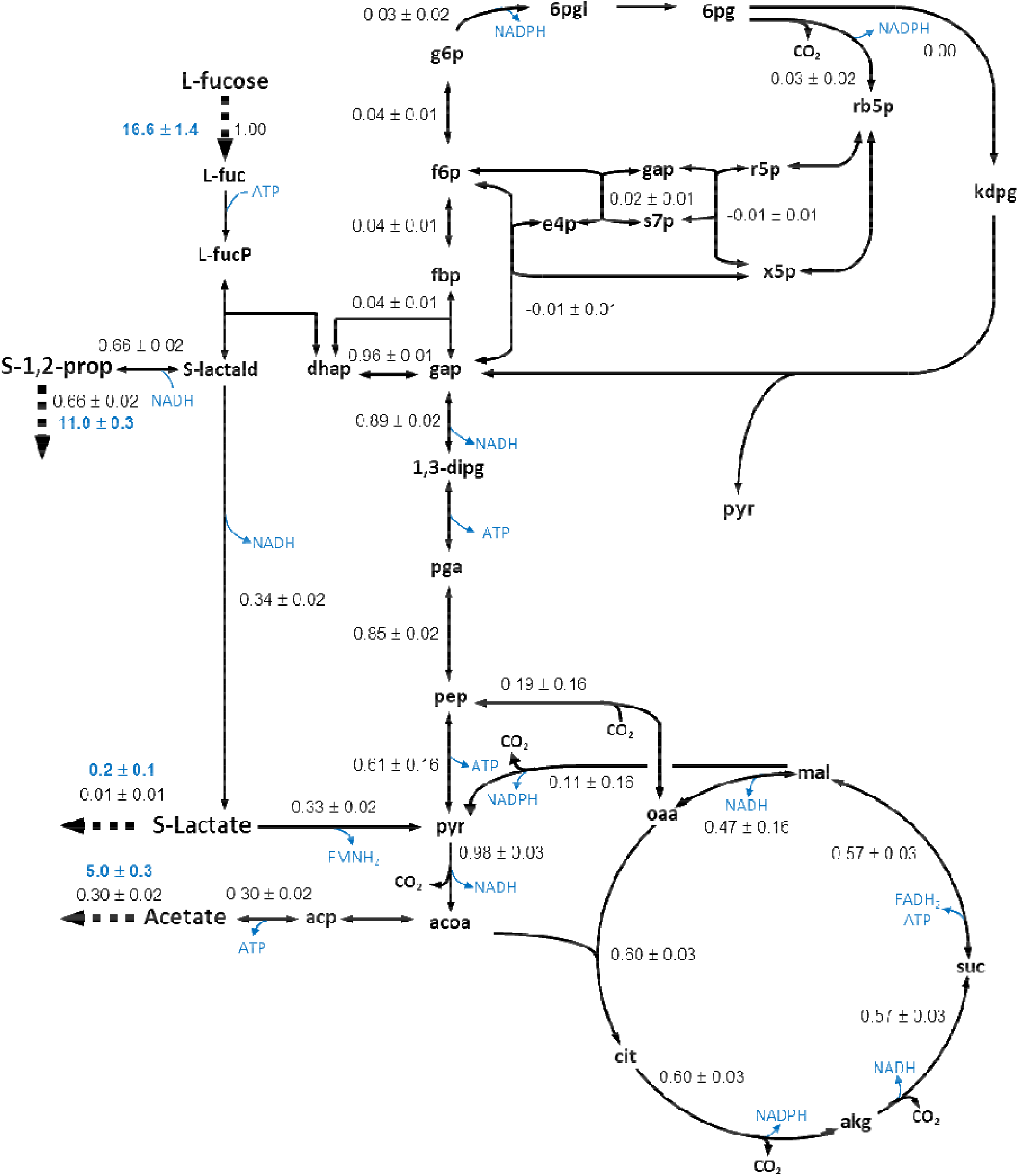
Flux distribution in the central metabolism of *Escherichia coli* K-12 MG1655 growing aerobically on fucose. Fluxes are given as a molar percentage of the specific fucose uptake rate which was set to 1. Net extracellular fluxes measured (mmol.(g_CDW_.h)^-1^) are shown in blue. The growth rate was 0.43 ± 0.04 h^-1^.

The flux distribution is mapped in Figure 4, with net fluxes expressed relative to the fucose uptake rate. Approximatively two-thirds of the S-lactaldehyde formed from the breakdown of fuculose-1-phosphate goes into the formation of 1,2-PDO, with the remaining flux being directed towards the pyruvate node through AldA. This is consistent with the fact that the lactate detected at trace concentrations in the culture supernatant (excretion flux ∼0.2 ± 0.1 mmol.(g_CDW_.h)^-1^) was found to be the S enantiomer (Table S1). Furthermore, DHAP is overwhelmingly (∼90%) channelled into the endergonic part of glycolysis, with the flux through the pentose phosphate pathway reduced to the minimum required to satisfy anabolic demand for carbon precursors (ribose-5-phosphate and erythrose-4-phosphate). In line with previous results (24), the flux through the Entner-Doudoroff pathway was zero. The carbon flux merging at the pyruvate node is thus massive in spite of the amounts wasted in the excretion of 1,2-PDO. For example, the total absolute flux through the pyruvate dehydrogenase complex is twice as high on fucose (16.3 ± 0.4 mmol.(g_CDW_.h)^-1^) as on glucose under similar conditions (8.58 ± 0.7 mmol.(g_CDW_.h)^-1^) (25)). Although the acetyl-coA produced by the pyruvate dehydrogenase complex is mostly directed towards the tricarboxylic acid cycle (60%), a significant fraction is channelled into acetate biosynthesis (30%). Acetate overflow is thus about 30% higher than commonly reported in the literature for growth on glucose (25).

The fluxes obtained can also be used to estimate the redox and energy metabolism of *E. coli* during aerobic growth on fucose. First, it turns out that NADPH supply from NADPH-dependent fluxes is higher than biomass-related anabolic NADPH requirements (Figure 5A). On fucose, NADPH is mainly (∼80%) supplied by isocitrate dehydrogenase, covering 140% of the anabolic demand, whereas on other substrates such as glucose, NADPH is supplied mainly by the pentose phosphate pathway (25). The apparent excess of NADPH may be converted into NADH through the activities of the soluble and membrane-bound, proton-translocating transhydrogenases UdhA and PntAB (26).

**Figure 5.**
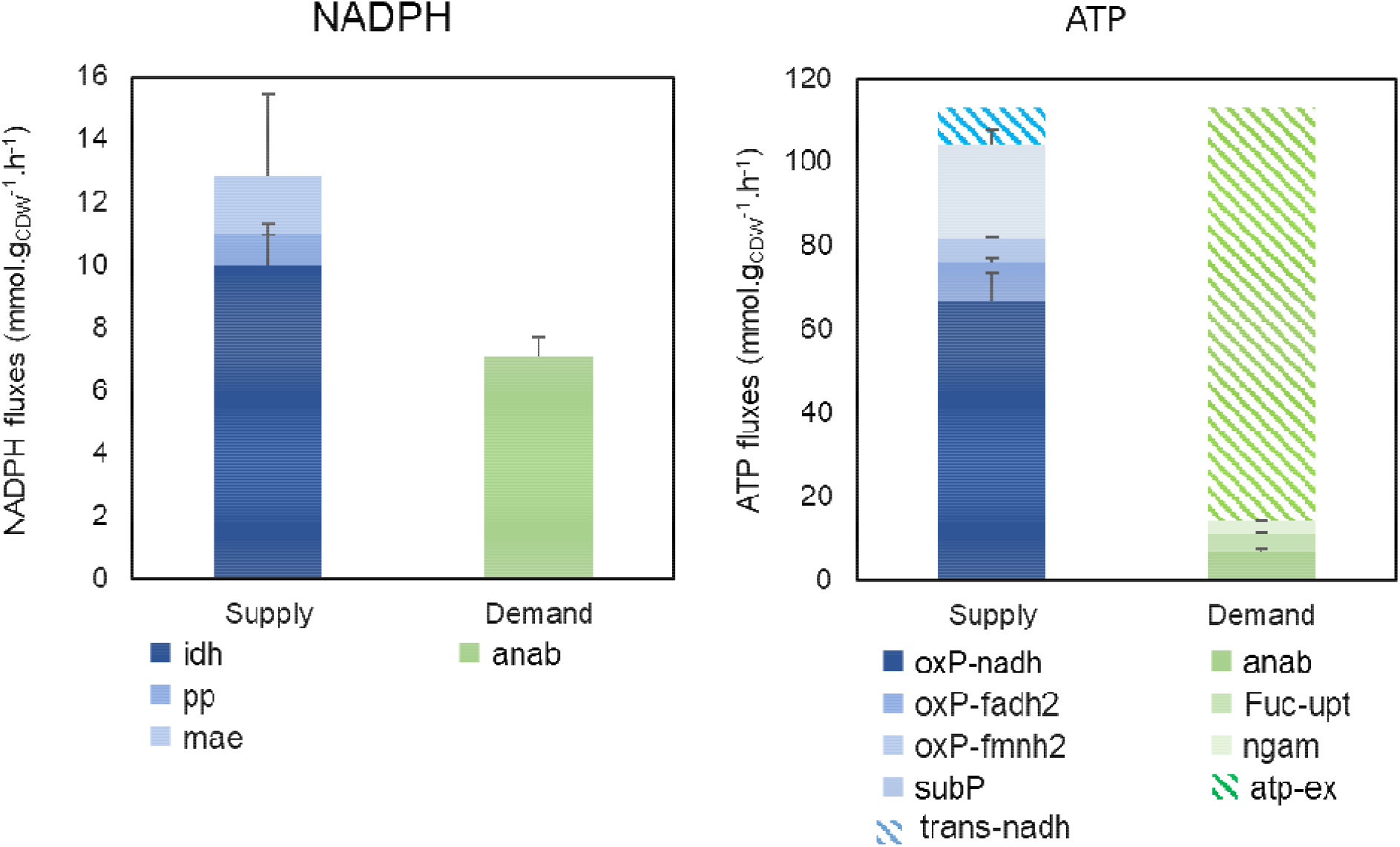
Quantitative analysis of redox and energy metabolism in *Escherichia coli* K-12 MG1655 grown aerobically on fucose. The absolute fluxes (mmol.(g_CDW_.h)^−1^) of reactions linked to NADPH (A) and ATP (B) metabolism are shown, as calculated from carbon fluxes (Figure 4).

For energy metabolism, ATP is supplied by substrate-level phosphorylation *i*.*e*. the difference between ATP-producing and ATP-consuming fluxes and from oxidative phosphorylation, considering NADH-, FADH_2_- and FMNH_2_-dependent fluxes and the conversion of these reducing equivalents into ATP (P/O ratio) (Figure 5B). The ATP demand is obtained by adding up the requirements for anabolism, non-growth-associated maintenance (27) and fucose uptake. The main source of ATP is NADH oxidation and to a lesser extend other redox equivalents; the contribution of substrate-level phosphorylation is minor (Figure 5B). Since the flux of reducing equivalents consumed in the reduction of S-lactaldehyde into 1,2-PDO is almost fully compensated by the flux of reducing equivalents produced by its oxidation into pyruvate, this indicates that 1,2-PDO does not contribute significantly to energy metabolism. Finally, ATP is utilized in roughly equal parts in fucose transport, non-growth-associated maintenance and anabolism. Together, these results indicate that *E. coli* K-12 generates a large apparent excess of ATP. This excess is available to cover the ATP required for growth-associated maintenance, which represents a very large part of cellular energy requirements (27). Indeed, growth-associated maintenance includes the unidentified and/or unquantifiable energetic requirements of macromolecule polymerization, gradient maintenance, protein folding, *etc*. (28), is itself difficult to quantify (27) and can vary substantially depending on growth conditions (29). It can also account for underestimated energy costs, such as that of fucose transport into cells, for which the amount of energy required is not yet well established.

### 6-deoxyhexose sugar metabolism in *E. coli* Nissle 1917

Under anaerobic conditions, the growth of *E. coli* Nissle 1917 on L-fucose and on L-rhamnose is characterized by two phases, with specific rates that are initially high (given in Table 2) and then drastically reduced (Figure S3). At this stage, we have no clear explanation for this phenomenon, which was not observed in *E. coli* K-12. *E. coli* Nissle 1917 seems to grow slightly faster on L-fucose than *E. coli* K-12, with no marked effect on the molar yields of by-products. On L-rhamnose, the growth parameters of both strains are similar.

**Table 2:** Growth parameters of *E. coli* Nissle 1917 cultivated aerobically and anaerobically on L-fucose or on L-rhamnose as sole carbon sources. The data shown are the specific rates for growth (μ_max_), substrate uptake (q_S_), 1,2-propanediol formation (q_PDO_), acetate formation (q_Ace_), and formate formation (q_For_), calculated from the data in Figure S3, and the yields of biomass (Y_X/S_), 1,2-propanediol (Y_PDO/S_), acetate (Y_Ace/S_) and formate (Y_For/S_).

| Parameters | L-fucose |  | L-rhamnose |  |
| --- | --- | --- | --- | --- |
|  | -O<sub>2</sub> | +O<sub>2</sub> | -O<sub>2</sub> | +O<sub>2</sub> |
| $\mu$ (h<sup>-1</sup>) | 0.30 ± 0.01 | 0.48 ± 0.01 | 0.32 ± 0.01 | 0.28 ± 0.01 |
| $q_s$ (mmol. (g<sub>CDW</sub>.h)<sup>-1</sup>) | 20.8 ± 1.0 | 12.8 ± 0.3 | 20.8 ± 0.3 | 6.9 ± 0.2 |
| $q_{\text{PDO}}$ (mmol. (g<sub>CDW</sub>.h)<sup>-1</sup>) | 19.3 ± 1.44 | 11.1 ± 0.2 | 17.3 ± 0.2 | 5.5 ± 0.1 |
| $q_{\text{Ace}}$ (mmol. (g<sub>CDW</sub>.h)<sup>-1</sup>) | 14.7 ± 1.7 | 4.0 ± 0.2 | 13.4 ± 0.2 | 0.2 ± 0.1 |
| $q_{\text{For}}$ (mmol. (g<sub>CDW</sub>.h)<sup>-1</sup>) | 12.0 ± 0.6 | ND* | 10.2 ± 0.8 | ND* |
| $q_{\text{CO}_2}$ (mmol. (g<sub>CDW</sub>.h)<sup>-1</sup>) | NM** | 15.3 ± 2.4 | NM** | 12.3 ± 0.7 |
| $Y_{X/S}$ (g<sub>X</sub>.mol<sup>-1</sup>) | 14.3 ± 0.7 | 37.2 ± 1.1 | 15.9 ± 0.3 | 41.4 ± 1.3 |
| $Y_{\text{PDO}}$ (mol.mol<sup>-1</sup>) | 0.92 ± 0.08 | 0.86 ± 0.03 | 0.86 ± 0.02 | 0.79 ± 0.03 |
| $Y_{\text{Ace}}$ (mol.mol<sup>-1</sup>) | 0.71 ± 0.09 | 0.31 ± 0.02 | 0.67 ± 0.01 | 0.03 ± 0.01 |
| $Y_{\text{For}}$ (mol.mol<sup>-1</sup>) | 0.58 ± 0.04 | ND* | 0.51 ± 0.01 | ND* |
| $Y_{\text{CO}_2}$ (mol.mol<sup>-1</sup>) | NM* | 1.19 ± 0.2 | NM* | 1.79 ± 0.1 |

\*: Not determined
\*\*: Not measured

Under aerobic conditions, *E. coli* Nissle 1917 also produces large amounts of 1,2-PDO, with molar yields slightly higher even than those measured for *E. coli* K-12. Note that on L-rhamnose, the growth rate of *E. coli* Nissle 1917 is significantly lower than *E. coli* K-12’s. This is quite surprising since *E. coli* Nissle 1917 typically grows faster on carbon sources than *E. coli* K-12 (30). In keeping with the slower growth rate, acetate overflow is also drastically reduced on L-rhamnose (Table 2). Just like the K-12 strain, *E. coli* Nissle 1917 produces the S enantiomer of 1,2-PDO on L-fucose and L-rhamnose (Figure S1).

These results indicate that *E. coli* Nissle 1917 and *E. coli* K-12 have very similar metabolisms when grown anaerobically and aerobically on 6-deoxyhexose sugar as the sole carbon source. Both strains have 1,2-PDO as a major metabolic end-product, even in the presence of oxygen.

## DISCUSSION SECTION

In this study, we investigated the metabolism of *E. coli* on L-fucose or L-rhamnose, the two main 6-deoxyhexose sugars in the bacterium’s natural environment. Two strains of *E. coli*, the K-12 MG1665 model strain and the Nissle 1917 probiotic strain, were therefore grown anaerobically and aerobically on L-fucose or L-rhamnose in a bioreactor, and extracellular metabolites were identified and quantified throughout the culture period. Metabolic flux distributions were quantified based on those measurements for anaerobic growth and using additional isotopic data for aerobic growth on L-fucose. Metabolic flux analyses were then performed to obtain a detailed picture of the central metabolism on these carbon sources. Under anaerobic conditions, *E. coli*’s metabolism is highly constrained by the need to balance redox equivalent fluxes and the limited means to generate ATP. The ATP required to sustain cellular growth can be only provided through the synthesis of acetate from acetyl-CoA. Consequently, DHAP, one of the two cleavage products of L-fuculose-1-phosphate (on fucose) and L-rhamnulose-1-phosphate (on rhamnose), should mostly be used in the endergonic part of the Embden-Meyerhof-Parnas pathway. The reduction of S-lactaldehyde, the second cleavage product, into 1,2-PDO is thus necessary to reoxidize the NADH produced by glyceraldehyde-3-phosphate dehydrogenase. This is indeed what we observed on fucose and on rhamnose under anaerobic conditions, where the flux distribution between the two metabolic branches was close to 1:1. The slight discrepancy is due to biomass-related demand for carbon and NADPH, which is relatively low under anaerobic conditions, biomass yield accounting for just ∼5% Cmol/Cmol of the carbon consumed by the cells as deoxyhexose sugars.

Under anaerobic conditions, ignoring biomass production, the theoretical maximum ATP yield is 1 mole per mole of fucose (1 fucose + 1 ADP + 1 Pi → 1 1,2-PDO + 1 acetate + 1 formate + 1 ATP). This is half what it is on glucose (1 glucose + 2 ADP + 2 Pi → 2 lactate + 2 ATP). The carbon-yields on fucose determined here based on the flux map (Figure 3, 1 fucose → 0.95 1,2-PDO + 0.88 acetate + 0.95 formate) are close to the theoretical values. ATP production is thus scarcely enough to support anabolic demands and provide the maintenance energy required for growth. Remarkably, the fucose consumption rate is much higher (27.5 ± 0.7 mmol.(g_CDW_.h)^-1^ – Table 1) than estimated on glucose under anaerobic conditions (17.8 ± 0.3 mmol.(g_CDW_.h)^-1^ (31)), and measured on fucose (15.3 ± 0.3 mmol.(g_CDW_.h)^-1^ – Table 1), glucose (7.7 ± 0.4 mmol.(g_CDW_.h)^-1^ (25)) and gluconate (12.8 ± 0.3 mmol.(g_CDW_.h)^-1^ – Figure S4) under aerobic conditions. From a physiological perspective, faster substrate utilization increases fluxes through the central metabolism and is a means for anaerobically growing cells to compensate for the apparently lower energy yields relative to respiratory metabolism (32).

Interestingly, we found that S-1,2-PDO is also excreted in large amounts (> 0.7 mole per mole of 6-deoxyhexose sugar consumed – Table 1 &2) when the cells are cultivated aerobically. This was not expected since it is assumed in the literature that S-lactaldehyde is oxidized to S-lactate by AldA rather than being reduced to S-1,2-PDO by POR, S-lactate then entering the central metabolism by being converted to pyruvate (11–13). This description of *E. coli*’s aerobic metabolism on deoxyhexose sugars is mainly based on the facts that i) AldA activity is only detected under aerobic conditions on fucose (13); ii) POR is partially post-transcriptionally inactivated under aerobic conditions (by about 70% (13)) while its expression is inducible both anaerobically and anaerobically by fucose (12); and iii) 1,2-PDO excretion would lead to an important overflow of reduced carbon, although reducing equivalents may in theory be re-oxidized by respiration. Our metabolic flux analysis shows that there is a significant flux through AldA (about 1/3 of the substrate uptake rate), in line with the fact that its activity has been detected by others when the cells are grown aerobically on fucose (11, 13). Regarding 1,2-PDO synthesis by POR, it is worth mentioning that significant 1,2-PDO production (approximatively 0.3 mole of 1,2-PDO per mole of fucose) under similar conditions has been reported previously in the literature (13) and that POR remains active, albeit partially (13, 33). However, our results indicate that residual POR activity appears to be sufficient to reduce S-lactaldehyde into 1,2-PDO. Moreover, despite 1,2-PDO overflow - and of acetate to a lesser extent, there is still a large excess of energy, suggesting that growth is not energetically constrained, with this surplus available to cover the demands of growth associated maintenance. Presumably also, the excess energy is crucial in allowing the cell to face various stresses and contributes to their survival and adaptation to environmental changes (3, 34).

S-lactaldehyde is at a metabolic node where it can be either oxidized by AldA or reduced by POR. It is therefore tempting to invoke the NADH/NAD^+^ ratio as a regulatory mechanism for the metabolic fate of S-lactaldehyde, especially since NADH is an inhibitor of AldA (11). In *Salmonella*, where AldA is completely absent, S-lactaldehyde is fully reduced to 1,2-PDO during aerobic growth on L-rhamnose (35). Moreover, S-lactaldehyde, and aldehydes in general (36), are assumed to be toxic compounds for *E. coli*’s growth (13, 37). 1,2-PDO overflow may thus be an efficient detoxification process that avoids intracellular accumulation of S-lactaldehyde.

Finally, our comparison of the laboratory model strain K-12 MG16655 with *E. coli* Nissle 1917, a non-pathogenic probiotic commensal strain known to be a good colonizer of the human gut (14), shows that their metabolisms on 6-deoxyhexose sugars are similar, with 1,2-PDO excretion observed in both strains. The 1,2-PDO yields are similar whether the cells are cultivated on fucose or rhamnose, under anaerobic or aerobic conditions. This suggests that 1,2-PDO should be present in *E. coli*’s vicinity and could serve as a substrate for other bacteria in the same ecological niche, including short chain fatty acid (SCFA) producers (38). It has very recently been shown that so-called cross-feeding based on 1,2-PDO (produced by rhamnose and fucose metabolization) influences the competitive fitness of other bacteria in the gut and is an important ecological process that shapes the gut microbiome and its metabolism (4).

## Materials and Methods

### Chemicals

1,2-Propanediol, S(L)-1,2-Propanediol, R(D)-1,2-Propanediol, fucose and rhamnose were purchased from Sigma-Aldrich Chemie (Saint-Quentin Fallavier, France). Propane-2-ol, hexane and ethyl acetate (both of HPLC-grade) were purchased from VWRProlabo (Fontenay Sous bois, France).

### Bacterial Strains and Cultures

The strains used in this work were *Escherichia coli* K-12 MG1655 and the probiotic *Escherichia coli* Nissle 1917 strain. *E. coli* strains were grown on minimal synthetic medium containing per litre: 10 g of L-fucose or L-rhamnose, 2 g KH_2_PO_4_, 0.5 g NaCl, 2 g NH_4_Cl, 2 mmol MgSO_4_, 0.04 mmol CaCl_2_, and 1 mL of a microelement solution containing 15g.L^-1^ Na_2_EDTA·2H_2_O, 4.5 g.L^-1^ ZnSO_4_·7H_2_O, 0.3 g.L^-1^ CoCl_2_·6H_2_O, 0.1 g.L^-1^ MnCl_2_·4H_2_O, 0.1 g.L^-1^ H_3_BO_3_, 0.04 g.L^-1^ Na_2_MoO_4_·2H_2_O, 0.3 g.L^-1^ FeSO_4_·7H_2_O, and 0.03 g.L^-1^ CuSO_4_·5H_2_O. The medium for *E. coli* K-12 MG1655 also contained 0.1 g.L^-1^ thiamine. Magnesium sulphate and calcium chloride were autoclaved separately. Solutions of 6-deoxyhexose (fucose or rhamnose), microelements and thiamine were sterilized by filtration. Exponentially growing cells pre-cultured in the same medium were harvested by centrifugation, washed in the same volume of fresh medium (lacking carbon sources and thiamine) and inoculated at 1% (v/v). Cultures were performed in parallel with both strains in 500 mL bioreactors (Multifors system, INFORS HT, Switzerland). The pH was maintained at 6.9 throughout the fermentation process by automatic addition of 2M NaOH and the temperature was regulated at 37°C.

For the anaerobic cultures, the medium was flushed with N_2_ before inoculation. An N_2_ atmosphere was maintained throughout the culture period by gently and aseptically bubbling N_2_. The stirring speed was 300 rpm in a working volume of 0.3 L. For the aerobic cultures, aeration and the speed of the stirrer were adjusted to maintain adequate aeration (dissolved oxygen tension - DOT > 30 % saturation), in a working volume of 0.3 L. The percentage concentrations of O_2_, CO_2_ and N_2_ were measured in the gas output during the culture process using a Dycor ProLine Process mass spectrometer (Ametek, Berwyn, PA, USA).

### Determination of biomass and extracellular metabolite concentrations

Bacterial growth was monitored spectrophotometrically at 600 nm (OD_600_) (Genesys 6, Thermo, Carlsbad, CA, USA). For cell dry weight (CDW) measurements, cells were collected by vacuum filtration on pre-weighed and dried membrane filters (0.2 μm, regenerated cellulose, 47 mm, Sartorius, France). The filters were then washed with 0.9% NaCl and dried at 60 °C under partial vacuum for at least 24 h until a constant weight was achieved. The OD-CDW correlation factors obtained were CDW [g.L^−1^]=0.49×OD_600_ for *E coli* K-12 MG1655 and CDW [g.L^−1^]=0.53×OD_600_ for *E coli* Nissle 1917. The carbon, hydrogen, oxygen and nitrogen contents of the biomass were determined by elemental analysis (Ecole de Chimie, Toulouse). The biomass formulas were C_1_H_1.783_N_0.241_O_0.445_ for *E. coli* K-12 MG1655 and C_1_H_1.764_N_0.242_O_0.442_ for *E coli* Nissle 1917.

Fucose, rhamnose, acetate, 1,2-propanediol and other fermentation products (lactate, formate, ethanol) were quantified by nuclear magnetic resonance (NMR) (39).

### ^*13*^C-labelling experiments

^13^C-labelling experiments were performed using a mixture of 20% (mol/mol) [U-^13^C] fucose and 80% (mol/mol) [1-^13^C] fucose (99% of ^13^C atom, Eurisotop, France). To minimize the amounts of labelled substrate used, the cultures were performed in 1L baffled shake-flasks containing 80 mL of medium. The inoculation procedure was the same as described above. Samples of intracellular metabolites were collected in mid-exponential phase to ensure they reflected isotopic and metabolic pseudo-steady states. Mass fractions of intracellular metabolites and of proteinogenic amino acids were determined in triplicate respectively by ion chromatography–mass spectroscopy (IC–MS) as described previously (22, 40) and by gas chromatography–mass spectroscopy (GC–MS) as described previously (41). The isotopologue distributions were obtained by correcting for naturally occurring isotopes using IsoCor (https://github.com/MetaSys-LISBP/IsoCor/) (42). *Enantiomeric analysis of 1,2-propanediol and lactate*: The enantiomeric purity of 1,2-propanediol was determined by HPLC using a 250 x 4.6 mm, 5 μm particle size Chiralpak AD column (Daicel Industries, Tokyo, Japan). The mobile phase was hexane:propane-2-ol (95:5 v/v). The flow rate of the mobile phase was 1.0 mLmin^-1^. The column temperature was maintained at 30 °C and compounds were detected by refractrometry. The injection volume was 20 μL.

1,2-propanediol was first extracted by solvent from samples collected at the end of cultures separated from the cells by 10 min centrifugation at 4°C and 11000 g. Samples of supernatant (20 mL) were mixed with 20 mL of ethyl acetate, the mixture was shaken, and the organic phase was recovered. This step was repeated three more times. Sodium sulphate (anhydrous) was added into the organic phase to eliminate traces of water. Filtered organic phase was then evaporated under vacuum using a rotary evaporator. The temperatures of the water bath and condenser were 30 °C and 1°C respectively; the pressure was maintained at 100 mBar. Once the organic solvent was fully removed, the extracted extracellular metabolites were dissolved before injection in 1 mL of the mobile phase used for chromatographic analysis.

R- and S-Lactate were quantified in culture supernatants using Biosentec R/S lactic acid enzymatic kits according to manufacturer instructions (Biosentec, Toulouse, France). *Calculation of extracellular fluxes:* Extracellular fluxes (i.e. deoxyhexose sugar uptake, PDO, lactate, and acetate production, and growth rates) were determined from the time course concentrations of biomass, substrates and products using PhysioFit v1.0.1 (https://github.com/MetaSys-LISBP/PhysioFit, (43)). Standard deviation on extracellular fluxes were determined using a Monte-Carlo approach with 500 iterations.

### Metabolic fluxes analyses

Metabolic flux distributions in the central metabolism were obtained with stoichiometric models under the assumption of a metabolic pseudo-steady state, based on the network topology shown in Figure 1 and using experimental input and output fluxes as constraints (*i*.*e*. extracellular sugar uptake and by-products excretion rates, and anabolic requirements of precursors to support growth (28)). Under anaerobic conditions, the inactive reactions were removed from the metabolic network (Figure 3) and additional constraints were imposed to balance redox fluxes (*i*.*e*. the fluxes of NADPH and NADH production and utilization). Under aerobic conditions, methylglyoxal metabolism and the glyoxalate pathway were not included in the stoichiometric model. Fluxes between the tricarboxylic acid cycle and the lower part of glycolysis were considered to be carried by phosphoenolpyruvate carboxylase, phosphoenolpyruvate carboxykinase and malic enzyme as suggested by several authors (25, 44). Furthermore, because i) the soluble and membrane-bound, proton-translocating transhydrogenases UdhA and PntAB are active (26) and, ii) redox equivalents can be reoxidized by the electron transport chain, no co-enzyme balancing constraints were imposed. Instead, the metabolic fluxes were resolved under these conditions, using a stable isotope (^13^C) labeling approach, by fitting additional isotopic balance equations to the ^13^C-labeling patterns of metabolites. All flux calculations were performed using the sotware influx_si v5.3.0 (23). A chi-square statistical test was used to assess the goodness-of-fit for each condition. Standard deviations on the fluxes were determined using a Monte-Carlo approach with 100 iterations.

Redox (NADH, NADPH, FADH_2_ and FMNH_2_) fluxes were estimated by summing the fluxes through all reactions producing or consuming the corresponding cofactors. Similarly, ATP production via substrate-level phosphorylation was calculated by summing the fluxes of ATP-producing reactions and subtracting fluxes of ATP-consuming reactions. ATP produced *via* oxidative phosphorylation was inferred from the production rates of NADH, FADH_2_ and FMNH_2_, assuming P/O ratios of 1.5 for NADH, and 1.0 for FADH_2_ and FMNH_2_ (45). The ATP consumed for fucose transport *via* L-fucose-H^+^ symport activity was estimated assuming a stoichiometry of 1 H^+^ per internalized fucose (46). Finally, NADPH and ATP requirements for biomass formation were estimated from the composition of the biomass and measured growth rates. The non-growth-associated maintenance value was taken from the genome-scale model of Feist et al. (27).

## Acknowledgements

The authors thank MetaToul (Metabolomics &Fluxomics Facilities, Toulouse, France, www.metatoul.fr) and its staff for technical support and access to the NMR facility. MetaToul is part of the French National Infrastructure for Metabolomics and Fluxomics (www.metabohub.fr), funded by the ANR (MetaboHUB-ANR-11-INBS-0010). The authors are also grateful to Christoph Bolten &Christoph Wittmann for their help with GC-MS analyses.

## REFRENCES

1. Jiang N, Dillon FM, Silva A, Gomez-Cano L, Grotewold E. 2021. Rhamnose in plants - from biosynthesis to diverse functions. Plant Science 302:110687.

2. Albermann C, Distler J, Piepersberg W. 2000. Preparative synthesis of GDP-β-L-fucose by recombinant enzymes from enterobacterial sources. Glycobiology 10:875–881.

3. Alteri CJ, Mobley HL. 2012. Escherichia coli physiology and metabolism dictates adaptation to diverse host microenvironments. Current Opinion in Microbiology 15:3–9.

4. Cheng CC, Duar RM, Lin X, Perez-Munoz ME, Tollenaar S, Oh J-H, van Pijkeren J-P, Li F, van Sinderen D, Gänzle MG, Walter J. 2020. Ecological Importance of Cross-Feeding of the Intermediate Metabolite 1,2-Propanediol between Bacterial Gut Symbionts. Appl Environ Microbiol 86.

5. Conway T, Cohen PS. 2015. Commensal and Pathogenic Escherichia coli Metabolism in the Gut. Microbiol Spectr 3.

6. Chang D-E, Smalley DJ, Tucker DL, Leatham MP, Norris WE, Stevenson SJ, Anderson AB, Grissom JE, Laux DC, Cohen PS, Conway T. 2004. Carbon nutrition of Escherichia coli in the mouse intestine. Proc Natl Acad Sci U S A 101:7427–7432.

7. Autieri SM, Lins JJ, Leatham MP, Laux DC, Conway T, Cohen PS. 2007. L-fucose stimulates utilization of D-ribose by Escherichia coli MG1655 DeltafucAO and E. coli Nissle 1917 DeltafucAO mutants in the mouse intestine and in M9 minimal medium. Infect Immun 75:5465–5475.

8. Cocks GT, Aguilar T, Lin EC. 1974. Evolution of L-1, 2-propanediol catabolism in Escherichia coli by recruitment of enzymes for L-fucose and L-lactate metabolism. J Bacteriol 118:83–88.

9. Chen YM, Tobin JF, Zhu Y, Schleif RF, Lin EC. 1987. Cross-induction of the L-fucose system by L-rhamnose in Escherichia coli. Journal of Bacteriology 169:3712–3719.

10. Vía P, Badía J, Baldomá L, Obradors N, Aguilar J 1996. Transcriptional regulation of the Escherichia coli rhaT gene. Microbiology 142:1833–1840.

11. Baldomà L, Aguilar J. 1988. Metabolism of L-fucose and L-rhamnose in Escherichia coli: aerobic-anaerobic regulation of L-lactaldehyde dissimilation. J Bacteriol 170:416–421.

12. Chen YM, Lin EC. 1984. Dual control of a common L-1,2-propanediol oxidoreductase by L-fucose and L-rhamnose in Escherichia coli. J Bacteriol 157:828–832.

13. Zhu Y, Lin EC. 1989. L-1,2-propanediol exits more rapidly than L-lactaldehyde from Escherichia coli. J Bacteriol 171:862–867.

14. Grozdanov L, Raasch C, Schulze J, Sonnenborn U, Gottschalk G, Hacker J, Dobrindt U. 2004. Analysis of the genome structure of the nonpathogenic probiotic Escherichia coli strain Nissle 1917. J Bacteriol 186:5432–5441.

15. Bailey JK, Pinyon JL, Anantham S, Hall RM. 2010. Distribution of Human Commensal Escherichia coli Phylogenetic Groups. Journal of Clinical Microbiology 48:3455–3456.

16. Hobman JL, Penn CW, Pallen MJ. 2007. Laboratory strains of Escherichia coli: model citizens or deceitful delinquents growing old disgracefully? Mol Microbiol 64:881–885.

17. Boronat A, Aguilar J. 1981. Metabolism of L-fucose and L-rhamnose in Escherichia coli: differences in induction of propanediol oxidoreductase. J Bacteriol 147:181–185.

18. Clark DP. 1989. The fermentation pathways of Escherichia coli. FEMS Microbiol Rev 5:223–234.

19. Holms H. 1996. Flux analysis and control of the central metabolic pathways in Escherichia coli. FEMS Microbiol Rev 19:85–116.

20. Millard P, Enjalbert B, Uttenweiler-Joseph S, Portais J-C, Létisse F. 2021. Control and regulation of acetate overflow in Escherichia coli. Elife 10.

21. Altaras NE, Cameron DC. 1999. Metabolic Engineering of a 1,2-Propanediol Pathway in Escherichia coli. Appl Environ Microbiol 65:1180–1185.

22. Millard P, Massou S, Wittmann C, Portais J-C, Létisse F. 2014. Sampling of intracellular metabolites for stationary and non-stationary (13)C metabolic flux analysis in Escherichia coli. Anal Biochem 465:38–49.

23. Sokol S, Millard P, Portais J-C. 2012. influx_s: increasing numerical stability and precision for metabolic flux analysis in isotope labelling experiments. Bioinformatics 28:687–693.

24. Leatham MP, Stevenson SJ, Gauger EJ, Krogfelt KA, Lins JJ, Haddock TL, Autieri SM, Conway T, Cohen PS. 2005. Mouse intestine selects nonmotile flhDC mutants of Escherichia coli MG1655 with increased colonizing ability and better utilization of carbon sources. Infect Immun 73:8039–8049.

25. Nicolas C, Kiefer P, Letisse F, Krömer J, Massou S, Soucaille P, Wittmann C, Lindley ND, Portais J-C. 2007. Response of the central metabolism of Escherichia coli to modified expression of the gene encoding the glucose-6-phosphate dehydrogenase. FEBS Lett 581:3771–3776.

26. Sauer U, Canonaco F, Heri S, Perrenoud A, Fischer E. 2004. The Soluble and Membrane-bound Transhydrogenases UdhA and PntAB Have Divergent Functions in NADPH Metabolism of Escherichia coli. Journal of Biological Chemistry 279:6613–6619.

27. Feist AM, Henry CS, Reed JL, Krummenacker M, Joyce AR, Karp PD, Broadbelt LJ, Hatzimanikatis V, Palsson BØ. 2007. A genome-scale metabolic reconstruction for Escherichia coli K-12 MG1655 that accounts for 1260 ORFs and thermodynamic information. Mol Syst Biol 3:121.

28. Neidhart FC, Ingraham JL, Schaechter M. 1990. Physiology of the bacterial cell: a molecular approach. Sunderland (MA): Sinauer Associates Inc.,U.S.

29. Cheng C, O’Brien EJ, McCloskey D, Utrilla J, Olson C, LaCroix RA, Sandberg TE, Feist AM, Palsson BO, King ZA. 2019. Laboratory evolution reveals a two-dimensional rate-yield tradeoff in microbial metabolism. PLOS Computational Biology 15:e1007066.

30. Revelles O, Millard P, Nougayrède J-P, Dobrindt U, Oswald E, Létisse F, Portais J-C. 2013. The carbon storage regulator (Csr) system exerts a nutrient-specific control over central metabolism in Escherichia coli strain Nissle 1917. PLoS One 8:e66386.

31. Schütze A, Benndorf D, Püttker S, Kohrs F, Bettenbrock K. 2020. The Impact of ackA, pta, and ackA-pta Mutations on Growth, Gene Expression and Protein Acetylation in Escherichia coli K-12. Front Microbiol 11.

32. Koebmann BJ, Westerhoff HV, Snoep JL, Nilsson D, Jensen PR. 2002. The glycolytic flux in Escherichia coli is controlled by the demand for ATP. J Bacteriol 184:3909–3916.

33. Cabiscol E, Hidalgo E, Badía J, Baldomá L, Ros J, Aguilar J. 1990. Oxygen regulation of L-1,2-propanediol oxidoreductase activity in Escherichia coli. J Bacteriol 172:5514–5515.

34. Jones SA, Chowdhury FZ, Fabich AJ, Anderson A, Schreiner DM, House AL, Autieri SM, Leatham MP, Lins JJ, Jorgensen M, Cohen PS, Conway T. 2007. Respiration of Escherichia coli in the Mouse Intestine. Infect Immun 75:4891–4899.

35. Baldoma L, Badia J, Obradors N, Aguilar J. 1988. Aerobic excretion of 1,2-propanediol by Salmonella typhimurium. Journal of Bacteriology 170:2884–2885.

36. Kunjapur AM, Prather KLJ. 2015. Microbial Engineering for Aldehyde Synthesis. Appl Environ Microbiol 81:1892–1901.

37. Subedi K, Kim I, Kim J, Min B, Park C. 2008. Role of GldA in dihydroxyacetone and methylglyoxal metabolism of Escherichia coli K12. FEMS microbiology letters

38. Louis P, Flint HJ. 2017. Formation of propionate and butyrate by the human colonic microbiota. Environmental Microbiology 19:29–41.

39. Enjalbert B, Letisse F, Portais J-C. 2013. Physiological and Molecular Timing of the Glucose to Acetate Transition in Escherichia coli. Metabolites 3:820–837.

40. Kiefer P, Nicolas C, Letisse F, Portais J-C. 2007. Determination of carbon labeling distribution of intracellular metabolites from single fragment ions by ion chromatography tandem mass spectrometry. Anal Biochem 360:182–188.

41. Wittmann C, Hans M, Heinzle E. 2002. In vivo analysis of intracellular amino acid labelings by GC/MS. Anal Biochem 307:379–382.

42. Millard P, Letisse F, Sokol S, Portais J-C. 2012. IsoCor: correcting MS data in isotope labeling experiments. Bioinformatics 28:1294–1296.

43. Peiro C, Millard P, de Simone A, Cahoreau E, Peyriga L, Enjalbert B, Heux S. 2019. Chemical and Metabolic Controls on Dihydroxyacetone Metabolism Lead to Suboptimal Growth of Escherichia coli. Appl Environ Microbiol 85.

44. Fischer E, Sauer U. 2003. Metabolic flux profiling of Escherichia coli mutants in central carbon metabolism using GC-MS. European Journal of Biochemistry 270:880–891.

45. Taymaz-Nikerel H, Borujeni AE, Verheijen PJT, Heijnen JJ, Gulik WM van. 2010. Genome-derived minimal metabolic models for Escherichia coli MG1655 with estimated in vivo respiratory ATP stoichiometry. Biotechnology and Bioengineering 107:369–381.

46. Gunn FJ, Tate CG, Henderson PJ. 1994. Identification of a novel sugar-H+ symport protein, FucP, for transport of L-fucose into Escherichia coli. Mol Microbiol 12:799–809

